# Streamlining the alignment of UAV and TLS forest point clouds: an approach based on ground and tree stems

**DOI:** 10.1101/2024.01.03.574086

**Authors:** Nicola Puletti, Simone Innocenti, Matteo Guasti

**Affiliations:** CREA, Research Centre for Forestry and Wood, Viale Santa Margherita 80-52100-Arezzo,Italy

**Keywords:** Forest structure, Terrestrial laser scanning, Unmanned Aerial System, Boreal forests, Stand architectural traits

## Abstract

Accurate co-registration of terrestrial and aerial point clouds may provide a high-resolution description of tree components for large forest areas. However, an automated approach for co-registering point clouds is still needed, given the challenges in geospatial data processing, particularly in complex topographical conditions. The main objective of this study is to present the application of a novel procedure for the co-registration of point clouds obtained from terrestrial and UAV surveys in Mediter-ranean forests.

## 1 Introduction

Recent developments in laser scanning technologies represent an opportunity that the forestry sector must begin to consider today (Beland et al. 2019). Laser scanning systems, adjustable to three platforms - terrestrial, aerial, and satellite - yield data with diverse resolutions, ranging from millimetric to centimetric. The prevalent laser scanning systems employed in forestry applications are Airborne Laser Scanning (ALS) for aerial surveillance and Terrestrial Laser Scanning (TLS) for ground-level observations. Airborne laser scanning (ALS) has been used in the forest environmental monitoring sector for many years for a variety of objectives: forest mensuration (Kankare et al. 2013; Alvites et al. 2022), forest ecology (Müller and Brandl 2009) water basin analysis (Bryndal and Kroczak 2019), road networks (Roussel et al. 2023) just to cite some main sectors. More recent, and limited, is the use of the terrestrial laser scanner (TLS), although the methodologies for analyzing TLS data are solid and efficient, both for forest mensuration (Calders et al. 2020) and forest structure monitoring (N. Puletti, Galluzzi, et al. 2021).

More recent is the use of laser scanning in forests with micro-lasers mounted on Unmanned Aerial Vehicles (UAVs) equipped with modern navigation receivers and real-time kinematic positioning (RTK) (Torresan et al. 2017). Such integrated systems, already known as UASs (Unmanned Aerial Systems), can considerably improve the accuracy of forest measurements. Compared with ALS data, UASs offer low material and operational costs along with high-intensity data collection (Giannetti et al. 2020). In addition, UAS missions can be planned flexibly, avoiding inadequate weather conditions, and providing data availability on-demand (Guimarães et al. 2020).

Given all these data sources, the last frontier for forest monitoring relates to the alignment of TLS, and ULS to enhance quantitative characterization of forest stands. Accurate tree heights are measured using LiDAR-UAS (i.e. ULS) returns and the tree positions and structure mainly based on TLS, so that aligning terrestrial and ULS scans results in an improvement of measurement accuracy (Giannetti et al. 2018). In summary, obtaining complete structural information is the main result of aligning point clouds from different data sources (Shao et al. 2022).

Efficient co-registration is the basis for studies oriented to coupling remote with proximal data. The currently available methods for co-registration can be classified into two categories: (1) transformation and registration and (2) feature matching. The first involves applying rigid or non-rigid transformations to align the point clouds with fixed georeferenced positions, which GPS devices can acquire. Rigid transformations include translation, rotation, and scaling, while non-rigid transformations may involve more complex deformations. Iterative methods, like the Iterative Closest Point (ICP), can optimize the alignment by minimizing the differences between corresponding points in the overlapping areas. Feature matching techniques aim to identify common features in overlapping areas of the point clouds. Within a forest, these features could be natural objects (e.g., big trees, rocks, or the ground) or artificial targets positioned strategically in the area of interest. Matching algorithms, such as ICP, can be employed to iteratively refine the alignment of the point clouds based on the identified common features (see for example (Nicola Puletti et al. 2022)).

This short communication presents preliminary results of a novel approach for TLS and ULS data co-registration, performed in three forest sites with different structural features. We mainly aim to define some practical and technical settings useful for further studies and analysis in the Mediterranean forest.

## 2 Material and methods

### 2.1 Data collection

#### 2.1.1 Study area

The study was carried out over three test sites of pure Beech (Fagus sylvatica), located in a mountain forest in Central Italy (Alpe di Catenaia, Arezzo). The climate at the study site was temperate, with warm, dry summers and cold, rainy winters. The mean annual rainfall was 1224 mm, and the mean annual temperature was 9.5°C. The stands showed differences in stand density, basal area and mean height (Table 1), while species composition is pure in all the test sites (Figure 1). The first test site considered in this experiment (F00) is a coppice abandoned from management (i.e. unthinned since 1972) with natural evolution patterns. Tree density is high here (about 2780 trees per hectare) due to the high number of shoots derived from previous coppice management. The other two test sites were established within managed high forest stands (conversion system with periodic thinning), one (F02) placed close to F00, and the other (F01) in the Southern part of the forest. F00 and F02 belong to an experimental trial established in 1972 (Chianucci et al. 2016).

**Table 1:** Tree site quantitative characteristics as obtained from TLS data collection. Those data were used as a reference in this experiment.

| id ads | N | mean dbh | sd of dbh | basal area |
| --- | --- | --- | --- | --- |
| F00 | 196 | 21 | 12.7 | 35.8 |
| F01 | 32 | 30 | 7.3 | 9.6 |
| F02 | 18 | 40 | 7.9 | 9.6 |

**Figure 1.**
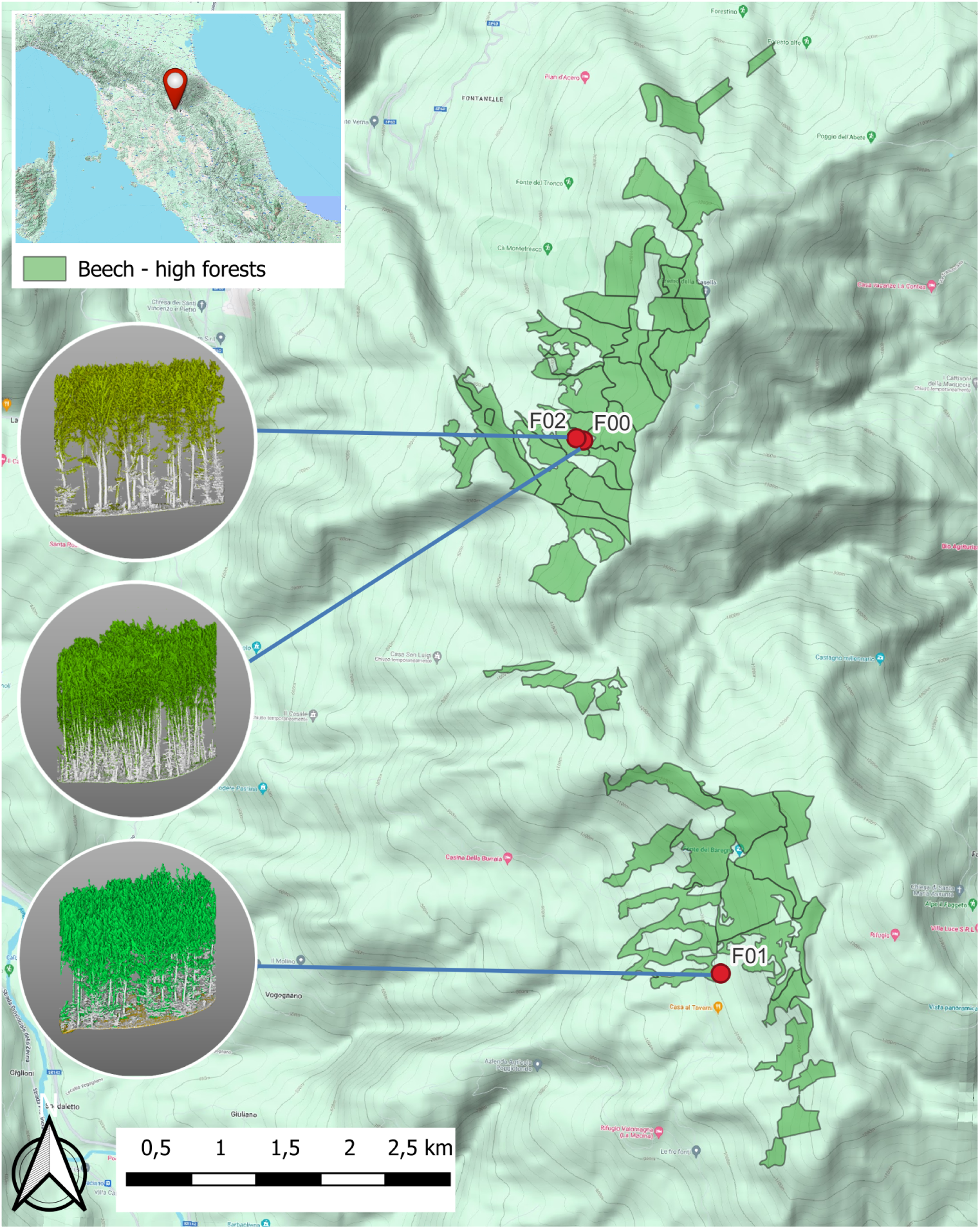
Geographic position of the three test sites. See Table 1 for quantitative details. One exemplative co-registered point cloud for each of the three test sites is represented on the circlesWhite points are from TLS, green points are from ULS.

#### 2.1.2 Field suveys

TLS data were collected in autumn 2023, during leaf off conditions, within circular plots of 15 m radius (707 m^2 of surface). The latitude and longitude of the plot centre were recorded by a nRTK receiver with positioning errors lower than 5 cm. TLS data were collected as described in a previous experience in Sila National Park (N. Puletti, Grotti, et al. 2021). The reported operative laser range outdoor is 15–20 m around the instrument.

In USL, different LiDAR camera settings were used over the same area. A total of 3 flights were performed for each test site during leaf-off conditions (autumn-winter 2023) with LiDAR camera set at -90°, -60°, and -45°.

### 2.2 Processing

#### 2.2.1 Alignment procedure

When downloaded from its data-logger, TLS data are centered on the starting point of the scan, which has relative coordinates x=0 and y=0. To align TLS to ULS point clouds, a translation was first carried out, adding to the TLS point cloud the coordinates of the center collected by the RTK GPS. Semi-automatic rotation on the Z axis was performed using Cloud Compare (http://cloudcompare.org) software by the *align* function (Figure 2). Trees and other objects (mainly the ground) identified by visual interpretation were used as corresponding points to stop the rotation process.

**Figure 2.**
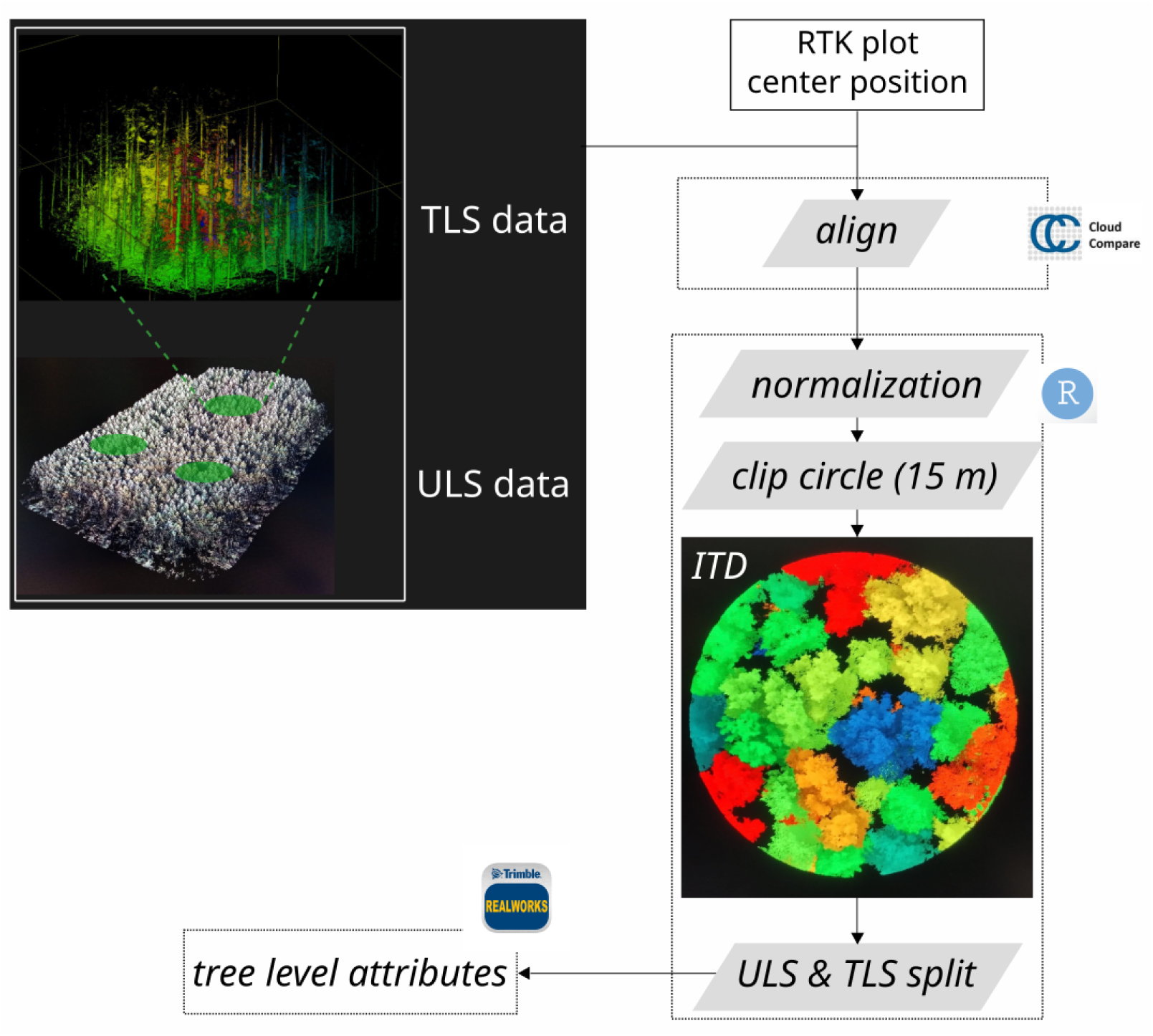
Workflow.

When merged, the resulting point cloud was normalized on the basis of ULS ground points classified by Terra®to a raster grid with a spatial resolution of 0.5 m.

#### 2.2.2 Tree attributes

From each TLS point cloud, for all living and dead trees with DBH >2.5 cm, we measured x and y position. Diameters were manually measured from TLS scans. After co-registration, from each ULS point clouds we estimated tree position using Individual Tree Detection (ITD) and Segmentation algorithms developed using the lidR package (Roussel et al. 2020) in *R* (Figure 2). We also repeated the same procedure to the same ULS-normalized point clouds filtered between 0 and 6 meters from the ground.

### 2.3 Relations between ULS and TLS data

The actual usability of aligned point clouds has been evaluated mainly using the position of the trees. Tree positions derived from aligned TLS data were used as reference to evaluate tree position derived by ULS. For such evaluation, we used Euclidean distance from the real position of the tree, measured by TLS, and the estimated positional value for the same tree, as obtained from ULS. We considered only trees within a radius of 15 m from the plot center.

## 3 Results

### 3.1 TLS data

From TLS data collection, we were able to determine the exact *xy* tree position with respect to the plot center (Figure 3). Table 1 summarizes the number of trees found, and other dimensional characteristics found in the three test sites. A total of 246 trees (196 in F00, 32 in F01, and 18 in F02) were found (Figure 3).

**Figure 3.**
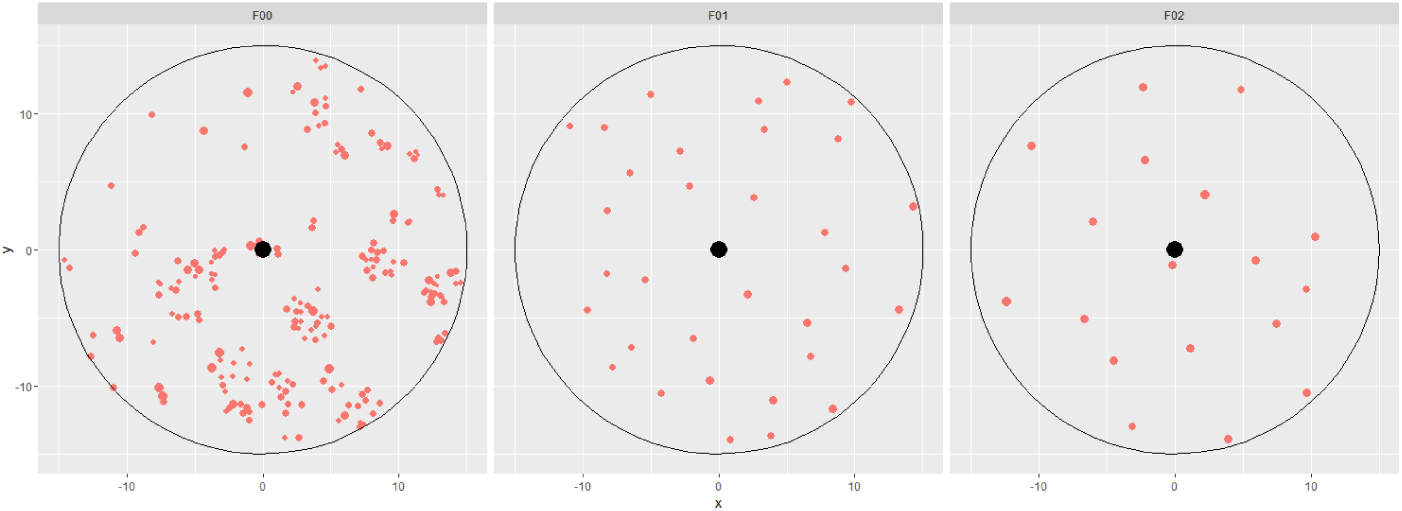
Position of the trees (orange circles) in the three test sites. Circles size is proportional to tree diameters. Black point is the plot center.

### 3.2 ULS data

For each test site we used different LiDAR camera angles -90° (nadir view), -60°, and - 45° as different flight missions. All the ULS point clouds always reach the ground with enough points to allow the creation of the digital terrain model (> 1500 ground points/m2). Nevertheless, the two no-nadir scans did not guarantee a good view of each tree from all directions, producing point clouds that are not useful for the subsequent alignment phase. For that reason, we prefer to not consider the scans with LiDAR camera set at -60° and -45° and use for alignment and further steps the nadir scan only.

From ULS_-90_, single trees can be easily detected, considering both leaf-off conditions and the really high point density: 4310 points m^−2^ for F00, 4858 points m^−2^ for F01, and 2698 points m^−2^ for F02.

Table 2 shows how Individual trees have been better detected using not the whole point clouds but ones normalized and cut at 6 meters from the ground. Figure 4 highlights how creating a cut off helps in ITD procedure.

**Figure 4.**
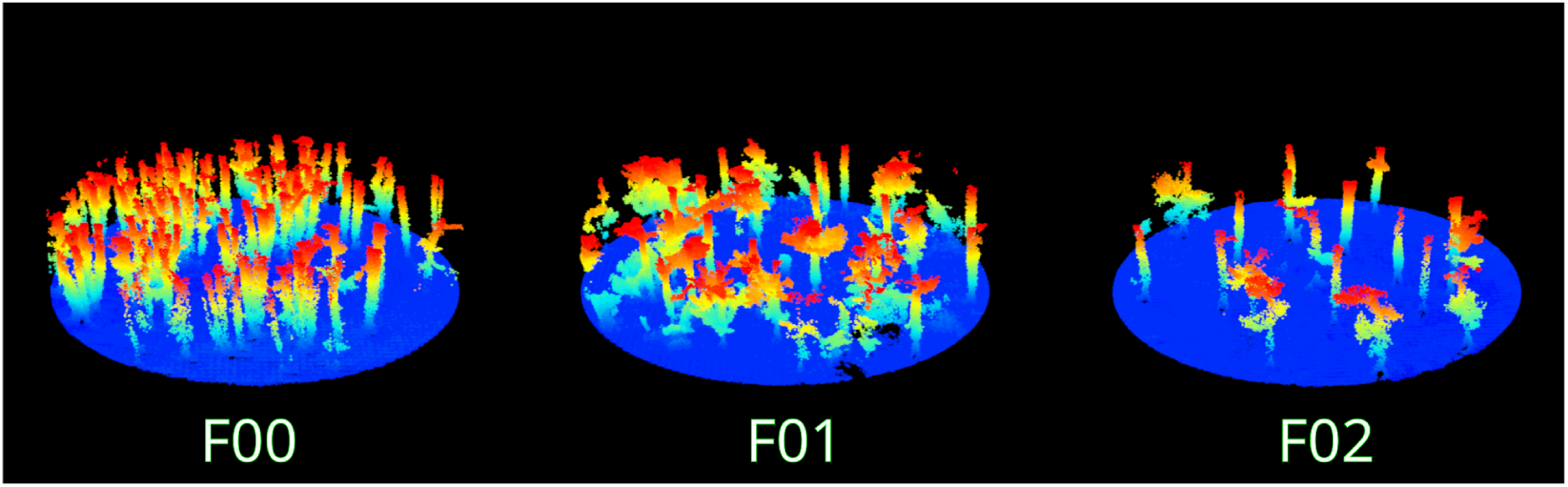
Three dimensional view of the point clouds normalized and cut at 6 m from the ground.

**Table 2:** Results on tree position detected from ULS data. Two types of point clouds (cut at 6m and whole) were assessed through RMSE (root mean squared error; m) and bias. Mean and SD were then derived.

| point cloud | id ads | obs trees (#) | detected trees (#) | RMSE of position | bias of position | SD of position |
| --- | --- | --- | --- | --- | --- | --- |
| whole stand | F00 | 196 | 29 | 1.71 | 1.55 | 0.72 |
| whole stand | F01 | 32 | 26 | 2.29 | 2.14 | 0.84 |
| whole stand | F02 | 18 | 23 | 3.30 | 2.77 | 1.85 |
| cut at 6 m | F00 | 196 | 36 | 1.58 | 1.33 | 0.87 |
| cut at 6 m | F01 | 32 | 32 | 2.03 | 1.41 | 1.49 |
| cut at 6 m | F02 | 18 | 18 | 1.08 | 0.80 | 0.75 |

RMSE and bias of tree positions as detected by ULS data. On everage, cut the normalized point clouds at 6 m from the ground performs better results.

## 4 Final remarks

- UAV technology is currently undergoing a dynamic development phase and holds the potential to offer foresters and researchers a portable remote sensing device suitable for real-time applications. This technology provides cost-effective options for collecting high-precision 3D data precisely for large forest covers when and where it is needed.
- The higher scanning density guaranteed by ULS flights allows a more detailed measure of the woody component of forest stands. In this experiment, just one flight with camera view set at -90° is enough. However, as for ALS scans and independent from forest structure, as well as for ULS, canopy leaves strongly limit the penetration of laser beams. When dimensional attributes of the trees are the main target, data collection should be carried out during leaf-off conditions for both TLS and ULS to ensure good results. Limiting the analysis to a reduced point cloud up to 6 m from the ground should be an efficient and effective option.
- Using leaf-off scans becomes essential to identify and position individual trees in broadleaved forests. It is known indeed that if an ITD algorithm that performs well across varying forest structures is still an open issue (Apostol et al. 2020), in the case of broadleaved species like beech, the available automatic methods are all barely suitable, offering insufficient tree detection rate. Broadleaved have more rounded crown shapes than coniferous trees, and the crowns tend to overlap near the top of the tree (Strunk and McGaughey 2023). The detection rate of individual trees, performed in
- our study over beech trees, was 100% in two forest structures (F01 and F02) over three, demonstrating the potential of the proposed methodology for broadleaved species. The third forest structure (F00) has many coppice shoots starting from the same stump, summing up in an intricate single crown, hardly detectable and separable from the others, for both ULS and TLS point clouds.
- Similar to previous findings, leaf-off scanning facilitates alignment procedures of ULS with TLS scans, as many details related to the ground and other reference objects can be better detected in both scans. Once aligned, the integrated TLS and ALS point cloud enhances the three-dimensional representation of the surveyed area.
- This experiment has considered three different LiDAR camera angles: -90°, -60°, and -45°. Integrating different scanning angles increases the density of points (i.e., more details) and is not very time expensive in the post-processing phase with the Terra® software. On the other hand, increasing the number of camera angles raises the number of field missions, with a noticeable impact on the time spent on the field and the need for charged batteries. From our experience, the nadir view (−90) is enough regarding costs and benefits.
- Once the scan area has been determined, the altitude and flight speed influence the mission duration and the point density together with the field of view (FOV) of the LiDAR camera. The FOV represents the angular extent of the laser beam or the scanner’s viewing angle, and it directly influences several aspects of LiDAR data acquisition. For example, a wider FOV allows the LiDAR sensor to capture a larger area in a single scan, leading to increased coverage. This can result in a higher point cloud density, especially in flat and open terrains. However, a narrow FOV may be preferred in complex terrains or areas with dense vegetation to focus the laser pulses more effectively. We verified that a flight altitude between 45 and 60 meters from the ground (depending on tree heights) and a flight speed between 3 and 5 m/s are useful settings to obtain the results presented in this study.

